# Modafinil Modulation of Oscillatory and Aperiodic Brain Activity in Rats Across Wakefulness and Subsequent Sleep

**DOI:** 10.64898/2026.01.14.699514

**Authors:** Mateo Mendoza, Diego M. Mateos, Juan Pedro Castro, Verónica Bisagno, Francisco J. Urbano, Pablo Torterolo, Alicia Costa

**Author notes:** Corresponding author: Alicia Costa.

## Abstract

Cortical states shape sensory processing, decision-making, and behavior, and are characterized by distinct patterns of neuronal oscillatory activity. Pharmacological agents can modulate these states and their underlying electrophysiological signatures. Modafinil is a widely used wake-promoting agent and cognitive enhancer, but its effects on cortical oscillations are not fully understood. In this study, we evaluated the effect of modafinil on cortical activity during wakefulness and subsequent sleep. Ten male Wistar rats were implanted with electrocorticographic electrodes and recorded following oral administration of modafinil or its vehicle. Our results show that modafinil increased wakefulness in a dose-dependent manner, with doses of 200 and 300 mg/kg producing prolonged wake episodes and reducing hypnogram complexity. During wakefulness, modafinil enhanced gamma-band power in anterior cortices, an effect primarily driven by increases in the aperiodic rather than oscillatory components. Although no sleep rebound was observed, subsequent NREM sleep exhibited increased delta power, particularly in anterior regions, indicating heightened sleep pressure. In conclusion, our findings demonstrate that modafinil-induced wakefulness is associated with an enhanced aperiodic gamma band component at the anterior cortices in the EEG. Furthermore, modafinil modulates sleep homeostasis by enhancing NREM sleep intensity rather than extending its duration.

## 1. INTRODUCTION

During wakefulness and sleep, the brain operates in distinct functional states that emerge from coordinated activity across cortical and thalamo-cortical networks, shaping sensory processing, cognition, and behavior (Harris and Thiele, 2011). These states are dynamically regulated by behavioral engagement and neuromodulatory systems, and are characterized by specific patterns of neuronal activity and oscillatory rhythms (Harris and Thiele, 2011; Zagha and McCormick, 2014). In rodents, the electroencephalogram (EEG) during wakefulness (W) is characterized by theta rhythms (4–9 Hz) concomitantly with gamma oscillations (30–100 Hz) (Adamantidis, Gutierrez Herrera and Gent, 2019). Gamma oscillations are thought to support brain processes that require distributed neuronal networks to communicate and integrate information. They have been associated with several cognitive processes such as perception (Gonzalez, Torterolo and Tort, 2023), attention (Engel, Fries and Singer, 2001), working memory (Lundqvist *et al*., 2016), and object representation (Tallon-Baudry and Bertrand, 1999). However, the interpretation of gamma activity as a dedicated mechanism of cortical processing has been challenged, emphasizing substantial variability in its origins and functional relevance (Ray and Maunsell, 2015). At a circuit level, a recent study has shown that gamma waves are triggered through recurrent connections between local excitatory and inhibitory neuronal populations that locally segregate neuronal assemblies through a winner-take-all computation, critical for cognitive processes (Gonzalez, Torterolo and Tort, 2023). In this regard, neuronal populations that generate gamma oscillations are targeted by various pharmacological agents that modulate gamma activity (Uhlhaas and Singer, 2010; González *et al*., 2021; Gallo *et al*., 2024; Castro-Zaballa *et al*., 2025).

Modafinil is a wake-promoting agent (or eugeroic drug) originally developed for the treatment of narcolepsy, but nowadays it is also indicated for other medical conditions with excessive somnolence (Nishino and Mignot, 2017; Billiard and Broughton, 2018). Modafinil has also been shown to improve performance in cognitive tasks in rodents (Morgan *et al*., 2007), healthy humans (Turner *et al*., 2003), and patients with neuropsychiatric disorders (Turner *et al*., 2004; Turner, 2006). This has led to its proposed use as a cognitive enhancer or nootropic drug (Battleday and Brem, 2015). However, despite these findings, its mechanism of action is not yet fully elucidated, possibly due to its complex pharmacological profile (Minzenberg and Carter, 2008). It is known that modafinil requires the dopamine transporter (DAT) for its effects (Wisor *et al*., 2001), and it may act through dopaminergic pathways by increasing extracellular dopamine levels by inhibition of the dopamine reuptake (Minzenberg and Carter, 2008). In vitro studies show that modafinil increases gap junction coupling, driving high-frequency coherence at the pedunculopontine and thalamocortical system, which my contribute to the generation of W (Garcia-Rill *et al*., 2007; Urbano, Leznik and Llinás, 2007). At the behavioral level, as a wake-promoting agent, modafinil increases time spent in wakefulness in both animals (Lin *et al*., 1992; Edgar and Seidel, 1997) and humans (Chapotot *et al*., 2003). Interestingly, modafinil does not induce a compensatory increase in sleep rebound, as measured by the duration of subsequent sleep episodes (Touret, Sallanon-Moulin and Jouvet, 1995), which suggests a complex interaction with sleep homeostasis. Although modafinil has been shown to reduce delta activity and increase high-frequency activity in the EEG of humans and mice during W (Del Percio *et al*., 2019; Vas *et al*., 2020), the specific patterns of oscillatory modulation it induces, particularly in the gamma band, remain poorly characterized.

In the present study, we investigated the effect of modafinil on cortical oscillations during W and subsequent sleep. Specifically, we aimed to characterize the EEG spectral composition during modafinil-induced W, focusing on gamma oscillations in rats. Additionally, we examined how prolonged modafinil-induced wakefulness influences subsequent sleep EEG dynamics.

## 2. MATERIALS AND METHODS

### 2.1 Experimental Animals

Ten male Wistar rats (250–300 g) were used. The number of animals was selected based on previous studies (Cavelli *et al*., 2017; Mondino *et al*., 2019, 2024). All experimental procedures were conducted in agreement with the National Animal Care Law (#18611) and were approved by the Institutional Animal Care Committee (Comisión Honoraria de Experimentación Animal de la Universidad de la República, and the Ethics Committee of the Facultad de Medicina; Exp. N° 070151-000011-22). Animals were housed under a 12:12-hour light/dark cycle (lights on at 9:00 a.m.) with food and water provided *ad libitum*. The room temperature was maintained at approximately 22 ± 2 °C throughout the experiments.

### 2.2 Surgical Procedures

Rats were prepared for chronic polysomnographic recordings using established protocols from our laboratory (Castro-Nin *et al*., 2024; Serantes *et al*., 2025). Anesthesia was induced with a mixture of ketamine-xylazine (90 mg/kg and 5 mg/kg, i.p., respectively). Pre-surgical analgesia was provided with ketoprofen (2.5 mg/kg, s.c.) (Ferry, Gervasoni and Vogt, 2014). Rats were positioned in a stereotaxic frame, and the skull was exposed. To record the electrocorticogram (ECoG), five stainless-steel screw electrodes were implanted in the skull, with their tips contacting the dura mater. As shown in Figure 1a, the electrodes were positioned according to Paxinos and Watson atlas (Paxinos and Watson, 2013) over the following structures: right olfactory bulb (rOb: L +1.25 mm, AP +7.5 mm), right primary motor cortex (rM1: L +2.5 mm, AP +2.5 mm), right primary somatosensory cortex (rS1: L +2.5 mm, AP −2.5 mm), and right secondary visual cortex (rV2: L +2.5 mm, AP −7.5 mm). The reference electrode was located over the cerebellum (C: L: 0 mm, AP: −11 mm). For electromyographic recordings (EMG), two electrodes were inserted into the subcutaneous space above the neck muscles. All electrodes were soldered to a socket and fixed onto the skull with acrylic cement. At the end of surgical procedures, ketoprofen (2.5 mg/kg, s.c.), and ceftriaxone (60 mg/kg; i.p.) were administered once daily for 48 hs post-surgery (Ferry, Gervasoni and Vogt, 2014). Incision margins were kept clean, and a topical antibiotic (bacitracin) was applied. Following surgery, animals were housed individually in transparent cages, with food and water *ad libitum*, and adapted to the recording chamber for five days before the start of the polysomnographic recordings.

**Fig. 1.**
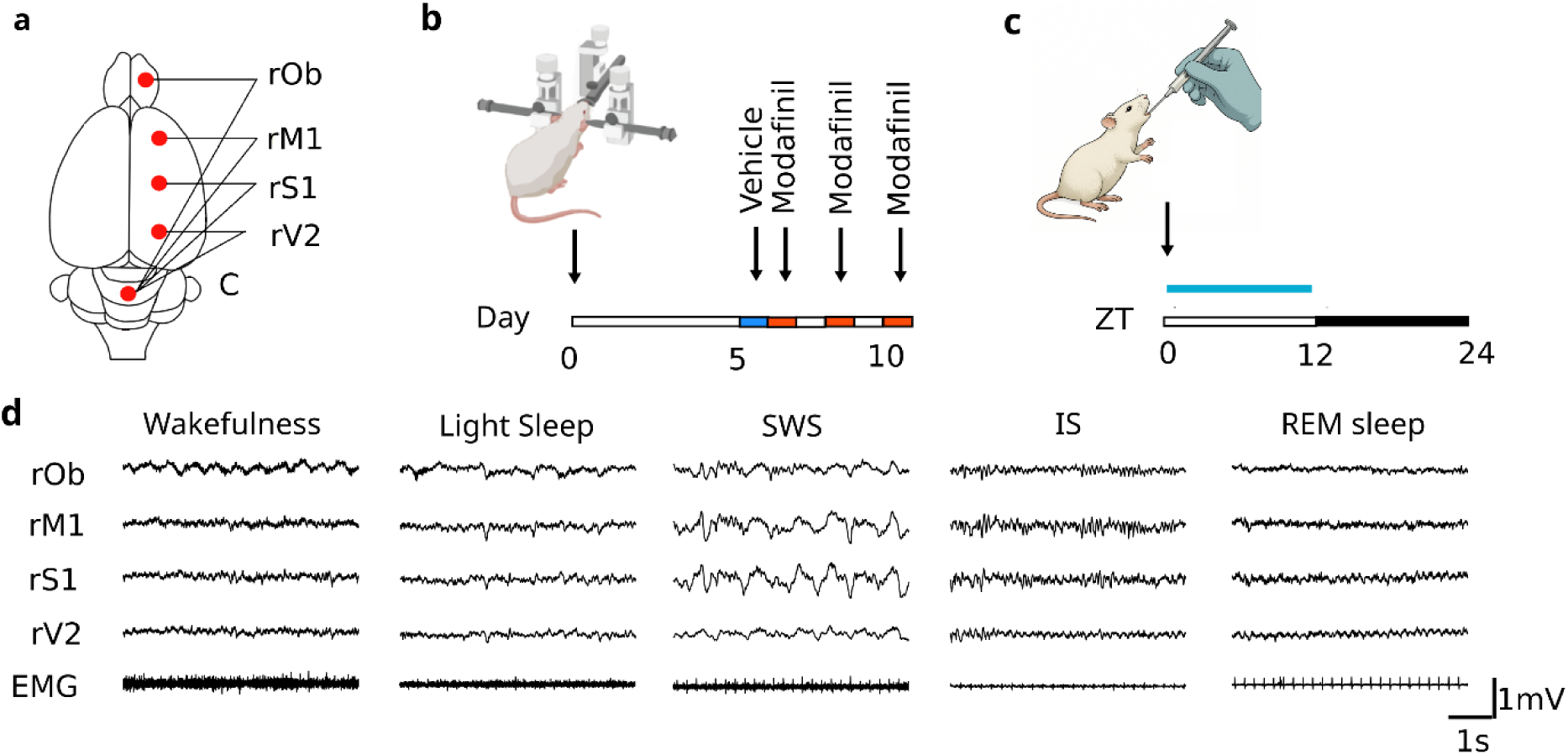
Experimental design and representative polysomnographic recordings. **(a)** Position of recording electrodes (red dots). The electrodes were referred to a common electrode located at the cerebellum (C), rOb: right olfactory bulb; rM1: right primary motor cortex; rS1: right primary somatosensory cortex; rV2: right secondary visual cortex. **(b)** Time course of the experimental protocol, numbers indicate days after surgery. **(c)** Timeline of a single experimental session, numbers indicate Zeitgeber Time (ZT). Rats (n=10) received vehicle or modafinil (100 mg/kg, 200 mg/kg, or 300 mg/kg) by oral gavage at ZT0 (arrow). The blue line represents the recording period. **(d)** Polysomnographic recordings of a representative animal after vehicle during the lights-on period. SWS: slow wave sleep, IS: intermediate state, EMG: electromyogram. Calibration bars indicate 1 second and 1 mV, respectively

### 2.3 Experimental sessions

#### 2.3.1 Administration of Modafinil

The experimental design is shown in Figure 1b and 1c. Modafinil (Activigil ®) was dissolved in distilled water and administered at three different doses: 100, 200, and 300 mg/kg, via oral gavage at the beginning of the light phase in a counterbalanced order. Vehicle (distilled water) was administered as control. Two to three days were left between sessions to avoid a possible cumulative effect. Because modafinil has low water solubility (Morgan *et al*., 2007), the suspension was freshly prepared before administration and thoroughly shaken to ensure a homogeneous distribution

#### 2.3.2 Polysomnographic recordings

Experimental sessions began at lights-on (9:00 am) and were conducted until 9:00 pm in a sound-attenuated chamber that also acts as a Faraday box. The polysomnographic recordings began immediately after the administration of modafinil and were performed through a rotating connector to allow the rats to move freely within the recording box. Bioelectric signals were amplified (×1000), filtered (0.1-500 Hz for ECoG and 10-500 Hz for EMG), sampled (1024 Hz, 16 bits), and stored in a PC (using Spike2 software version 9.04 (Cambridge Electronic Design, Cambridge, UK).

### 2.4 Data analysis

Data scoring was performed manually. The states of sleep and W were determined in 10-second epochs. W was defined as high-frequency, low-voltage activity in the ECoG and high EMG activity. Light sleep (LS) was characterized by a high-frequency, low-voltage ECoG background, occasionally interrupted by low-frequency (0.4-4 Hz), relatively high-amplitude waves, together with a lower level of EMG activity. Slow wave sleep (SWS) was defined as high-amplitude slow waves (0.4-4 Hz) and sleep spindles (12-14 Hz) primarily in M1 and S1 with reduced EMG activity (Torterolo *et al*., 2022). The intermediate state (IS) was defined as high-amplitude cortical spindles in anterior electrodes and theta rhythm (4-10 Hz) in the visual cortex with reduced EMG activity (Gottesmann, 1992; Serantes *et al*., 2025). REM sleep was defined as a predominant high-frequency and low-voltage activity in the ECoG of the anterior cortices, theta rhythm in the visual cortex, and marked reduction in EMG activity, except for occasional myoclonic twitches (Torterolo *et al*., 2022). See Figure 1d for examples of the recordings of these behavioral states.

Data analyses were performed using Python 3 as well as statsmodels (Seabold and Perktold, 2010), Scipy (Virtanen *et al*., 2020), and matplotlib libraries (Hunter, 2007). To quantify the temporal complexity of the sleep–wake architecture, we computed the Lempel–Ziv complexity (LZC) over the hypnogram, a metric based on pattern redundancy in symbolic time series (Lempel and Ziv, 1976). In the context of sleep staging, each sequential state in the hypnogram is treated as a symbol in the series, and LZC estimates the degree to which these state patterns are repetitive. LZC was computed over the first 6 hours of each experimental session, with values calculated per animal and condition. The specific LZC implementation used in this study is described in (Kaspar and Schuster, 1987). Power spectral density (PSD) was estimated using Welch’s method (SciPy; window = 1024 samples, overlap = None, fs = 1024 Hz, nfft = 10240), corresponding to a 1-second window, with a 0.5-second overlap and a frequency resolution of 0.1 Hz. Frequency bands were defined as delta (0.5–4 Hz), theta (4–10 Hz), sigma (10–16 Hz), beta (16–30 Hz), gamma (30–100 Hz), and high-frequency oscillations (HFO, 100–200 Hz). For W analyses, EEG segments corresponding to W epochs between 2–4 hours after vehicle or modafinil administration were concatenated, coinciding with the maximal wake-promoting effect of modafinil. To assess the effect of modafinil on subsequent sleep, PSDs were computed during SWS and REM sleep in the 7–9 hour post-administration window, when sleep time was maximal. To assess the temporal course of delta-band activity independently of sleep-wake state, delta power was extracted from spectrograms and normalized within each animal to its mean delta power across the entire recording. Normalized delta power was then averaged in hourly bins and across animals, allowing comparison of within-animal changes in low-frequency activity over time. Because EEG power spectra reflect a combination of periodic oscillatory activity and an aperiodic background component, spectral decomposition was performed using the Fitting Oscillations and One-Over-F (FOOOF) from FOOOF Python toolbox (Donoghue *et al*., 2020) as in a previous study in our group (Gallo *et al*., 2024). For this analysis, we focused on the power spectrum during W in the 1 and 150 Hz frequency range.

### 2.5 Statistics

Group data is shown as either mean ± SEM or regular boxplots showing the median, 1st, and 3rd quartiles. Data normality was verified through the Shapiro-Wilk test. For the contrast hypothesis, data were analyzed using a paired two-tailed t-test under conditions of normality; otherwise, a Wilcoxon signed-rank test was employed. To assess dose-related effects, a linear mixed-effects model (LMM) using the statsmodels package in Python was used. Doses (0, 100, 200, and 300 mg/kg) were included as a fixed effect, and subject (rat) as a random intercept. This approach accommodates the unbalanced design, as not all animals received all doses (n = 10 rats; 35 total observations). Model parameters were estimated using restricted maximum likelihood (REML), and the statistical significance of fixed effects was assessed using two-tailed z-tests (α = 0.05). For all analyses, statistical significance was set at p < 0.05

## 3. RESULTS

### 3.1 Modafinil increased wakefulness time and decreased the complexity of the sleep-wake cycle

To study the effect of modafinil, we begin by assessing its effect on sleep-wake architecture.

The hypnograms displayed in color code in Figure 2a show that modafinil administration (200 mg/kg) increased W and reduced sleep episodes. Modafinil produced a dose-dependent increase in the time spent in W during the first six hours after administration (Figure 2b, left; see also supplementary Figure 1). A LMM revealed that compared to vehicle modafinil at 200 mg/kg increased W time by 32.6% (*z* = 5.08, *p* < 0.001), while the 300 mg/kg dose increased it by 46.8% (*z* = 6.84, *p* < 0.001). In contrast, the 100 mg/kg dose did not differ significantly from vehicle (*p* = 0.544). Based on these results, the 200 mg/kg dose was selected for subsequent analyses as it elicited a significant wake-promoting response, avoiding potential ceiling effects. Accordingly, additional quantitative analyses of sleep–wake architecture during the first 6 h were performed for the vehicle and 200 mg/kg conditions. The mean duration and number of episodes of each behavioral state following 200 mg/kg of modafinil are exhibited in Table 1.

**Fig. 2.**
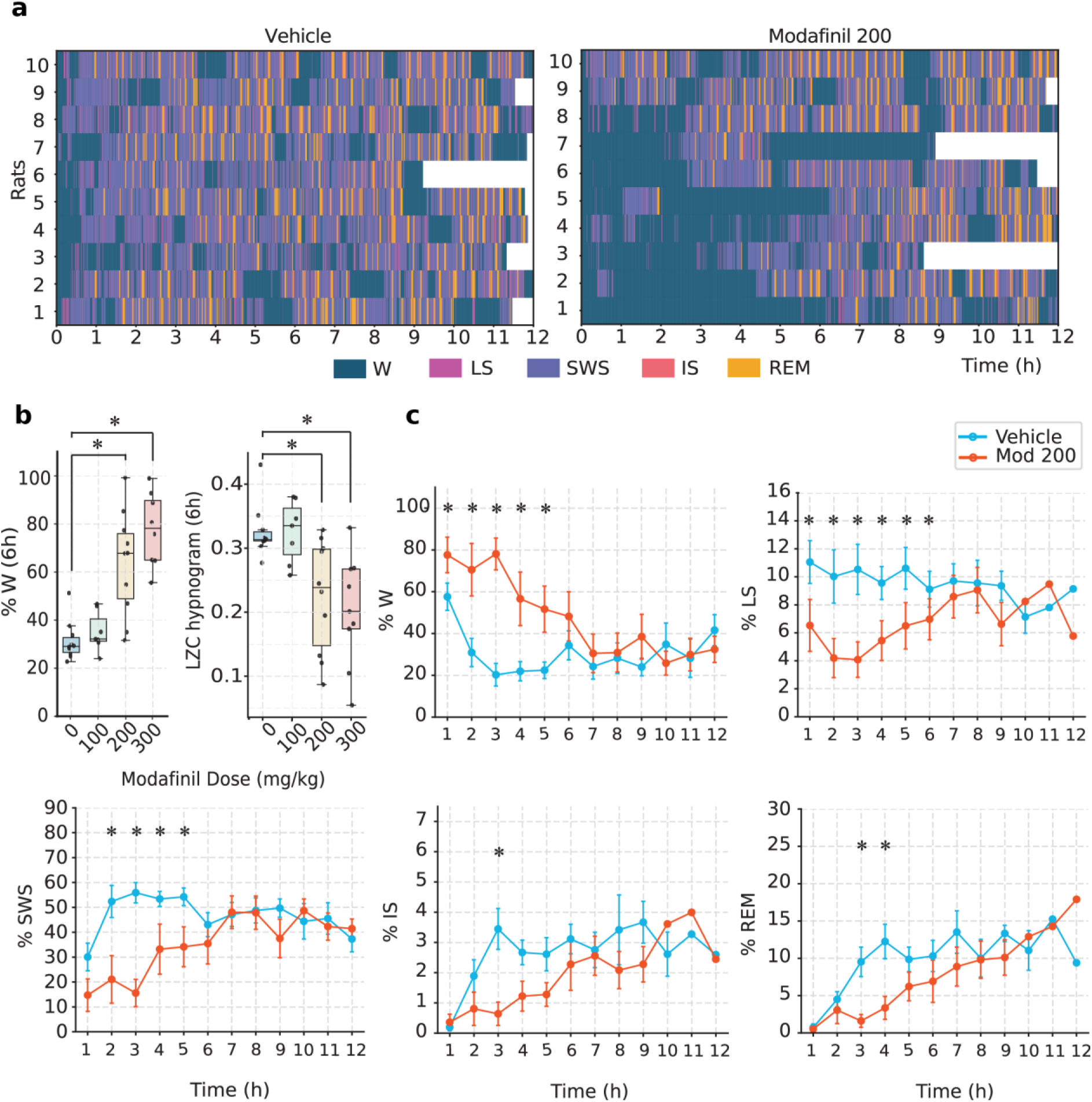
Effect of modafinil on the sleep-wake cycle. **(a)** Hypnograms in color code indicating sleep-wake stages under vehicle or modafinil 200 mg/kg (1 to 10 indicate the animaĺs number). **(b)** Left, percentage of time spent on wakefulness during the first six hours of treatment. Right, dose-dependent changes in the LZC of hypnograms over the first six hours of treatment. Statistical analysis was performed using LMM; significant differences from the vehicle are indicated by black lines with (*) (p < 0.05). **(c)** Time course of the percentage of time spent in each state per hour (mean ± SEM), comparing vehicle (light blue) and modafinil 200 mg/kg (orange). A paired t-test was used to assess differences between conditions; significant differences (p < 0.05) are indicated by (*). Sample size: n = 10 for hours 0 to 8; n = 7 for hours 9 to12. W: wakefulness, LS: light sleep, SWS: slow wave sleep, IS: intermediate state sleep, REM: REM sleep

**Table 1.** Comparative effects of vehicle vs. modafinil 200 mg/kg on sleep–wake architecture during the initial six-hour post-administration interval. Data are expressed as mean ± SEM. Normality was assessed using the Shapiro–Wilk test. Paired t-tests were applied when normality assumptions were met; otherwise, Wilcoxon signed-rank tests were used. P-values refer to the statistical significance of the difference between conditions (p < 0.05) and are indicated by (*)

|  | Vehicle | Modafinil 200 mg/kg | p-value |
| --- | --- | --- | --- |
| **Wakefulness** |  |  |  |
| Number of episodes | 96.4 ± 4.3 | 61.8 ± 12.0 | 0.03* |
| Mean duration (s) | 70.1 ± 4.3 | 705.3 ± 409.1 | <0.01* |
| **Light sleep** |  |  |  |
| Number of episodes | 128.4 ± 13.6 | 76.5 ± 15.2 | 0.01* |
| Mean duration (s) | 16.8 ± 0.7 | 18.6 ± 2.7 | 0.45 |
| **Slow wave sleep** |  |  |  |
| Number of episodes | 127.0 ± 10.6 | 71.3 ± 15.2 | <0.01* |
| Mean duration (s) | 86.2 ± 7.5 | 79.0 ± 12.1 | 0.59 |
| **Intermediate state** |  |  |  |
| Number of episodes | 25.1 ± 2.2 | 13.0 ± 4.0 | 0.02* |
| Mean duration (s) | 19.9 ± 0.8 | 15.7 ± 2.3 | 0.04* |
| **REM sleep** |  |  |  |
| Number of episodes | 23.0 ± 2.6 | 9.3 ± 2.7 | 0.01* |
| Mean duration (s) | 74.0 ± 6.1 | 78.9 ± 19.0 | 0.78 |

To evaluate the temporal structure of the sleep-wake cycle under modafinil, we computed the Lempel–Ziv complexity (LZC) of the hypnograms over the same first 6 hours of the light phase (Figure 2c, right). LZC values showed a baseline mean of 0.326 in the vehicle group (n = 10), with no apparent change at 100 mg/kg modafinil (mean = 0.332, n = 8). In contrast, modafinil at 200 mg/kg (mean = 0.232, n = 11) and 300 mg/kg (mean = 0.203, n = 9) produced marked decreases in LZC. Consistent with these observations, a LMM revealed a significant main effect of dose, with complexity reduced by 0.101 (z = –3.45, p = 0.001) and 0.122 (z = –4.06, p < 0.001) at 200 and 300 mg/kg, respectively, compared to vehicle. No significant difference was observed at 100 mg/kg (p = 0.918). These results indicate that modafinil increases wakefulness in a dose-dependent manner and reduces the complexity of the sleep-wake cycle, consistent with a shift toward a more predictable and less fragmented state. Finally, we analyzed the effect on the temporal distribution of sleep-wake states by performing an hour-by-hour comparison of the percentage of time allocated to each state under vehicle or modafinil at 200 mg/kg (Figure 2c). Hourly quantification of each state showed that modafinil significantly increased W, particularly during the first five hours post-treatment (p < 0.05, paired t-test). This increase was accompanied by a reduction in the time spent in SWS, LS, IS, and REM sleep. Notably, no rebound in the sleep time was observed after the modafinil wake-promoting effect, neither for 200 mg/kg nor 300 mg/kg (Supplementary Figure 1). In addition, modafinil increased the latency to REM sleep (mean ± SEM: 105.3 ± 15.4 vs 207.8 ± 41.5 min; p<0.05) and not to SWS (21.6 ± 3.8 vs 84.2 ± 35.8 min; p = 0.12) (see boxplot in Supplementary Figure 2).

### 3.2 Modafinil increased the gamma band spectral power of the anterior cortices

Next, we analyzed the ECoG power spectrum of W between the second and fourth-hour post-administration based on the results shown in Figure 2c. Figure 3a illustrates the effect of modafinil on the rM1 cortex in one animal. Figure 3b shows the mean ± SEM power spectral density (PSD) during W across cortical regions for all animals. Compared to W following vehicle administration, modafinil significantly increased gamma-band (30–100 Hz) power and HFO in the rOb, rM1, and rS1, while no significant effects were observed in rV2. Frequency bins with statistically significant differences (paired t-test, p < 0.05) are indicated by black dots at the upper section of Figure 3b plots.

**Fig. 3.**
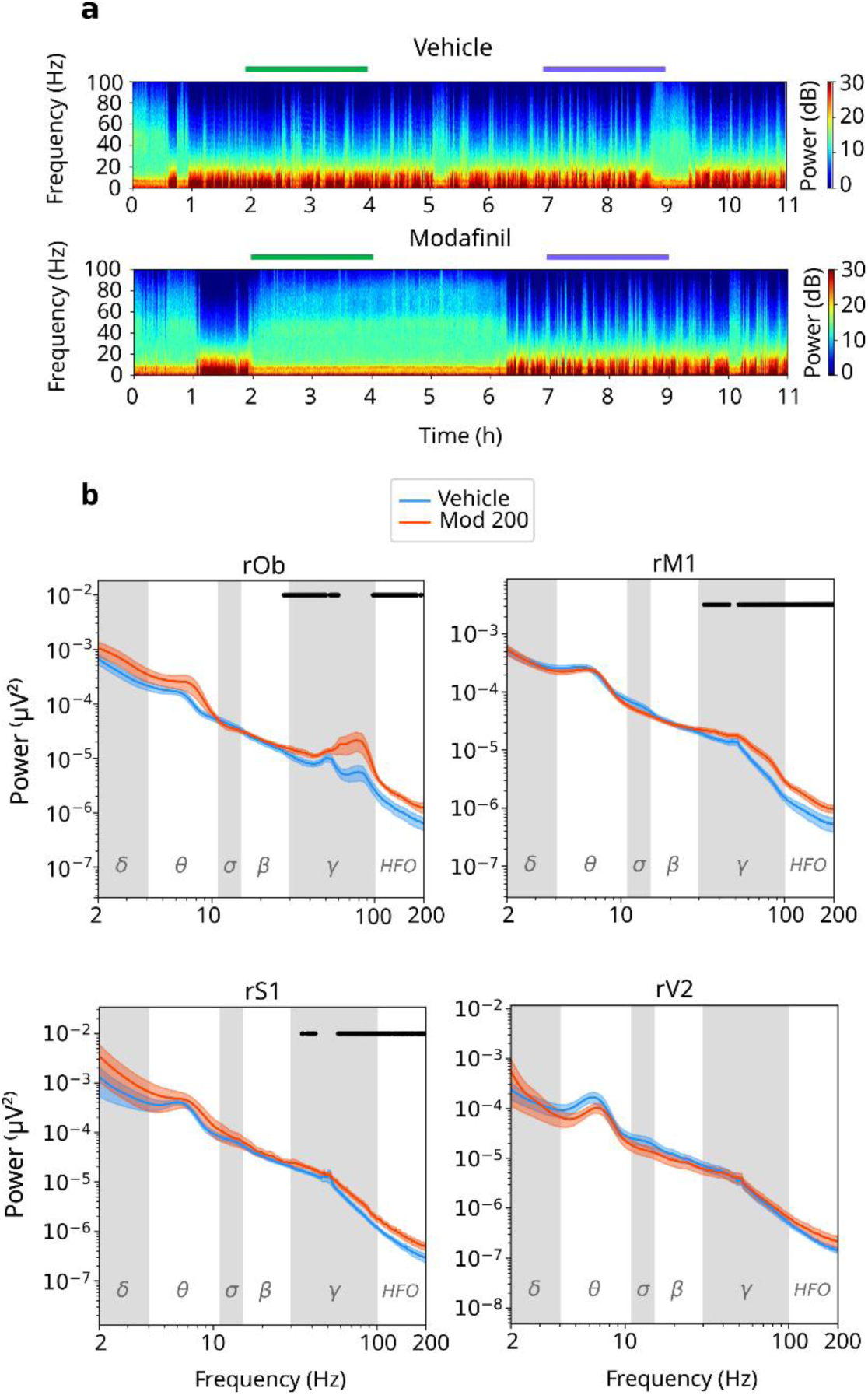
Effect of modafinil on wakefulness spectral power. **(a)** Representative spectrograms of rM1 in an animal over an 11-hour recording session following vehicle or modafinil (200mg/kg) administration. Lines above the spectrogram indicate analyzed time for wakefulness (green) and slow wave sleep or REM sleep (violet, see also Figure 5 and Supplementary Figure 3). **(b)** Power spectrum density during wakefulness from hours two to four post-administration of all the recorded cortices. Mean ± SEM. Background shading indicates canonical frequency bands: delta (0.5 - 4 Hz); theta (4 - 10 Hz); sigma (10 - 16 Hz); beta (16-30 Hz); gamma (30-100 Hz); and high-frequency oscillations (HFO, 100-200 Hz). Frequency bins showing significant differences between conditions (paired Student’s t-test, p < 0.05) are marked with black dots on upper sections of plots. rOb: right olfactory bulb (n = 6), rM1: right primary motor cortex (n=10), rS1, right primary somatosensory cortex (n=9), rV2: right secondary visual cortex (n=7)

The power spectrum can be described as comprising periodic components, reflected as narrowband oscillatory peaks, superimposed on an aperiodic background characterized by a 1/f-like decay (Voytek *et al*., 2015). Because increases in spectral power may arise from either enhanced oscillatory activity or changes in the aperiodic background signal (e.g., elevation in the spectral offset or slope flattening), we next applied the Fitting Oscillations and One-Over-F (FOOOF) algorithm (Donoghue *et al*., 2020) to decompose the PSD into its aperiodic (1/f-like) and periodic (oscillatory peak) components. Figure 4a shows a representative example of FOOOF model fitting applied to the mean PSD from the rM1, comparing vehicle and modafinil conditions. The panel displays from left to right, the total PSD, the full model, the aperiodic model, and the oscillatory model. Two oscillatory peaks were clearly identified, corresponding to theta and gamma bands. Since modafinil was described above to increase gamma-band PSD, we continue focusing our analysis on this frequency band. In fact, the increase in the gamma power under modafinil is closely related to the increase in the aperiodic signal. As shown in Figure 4b, both the aperiodic slope and offset were significantly altered by modafinil compared to the vehicle. Modafinil reduced the exponent values in rOb (1.43 ± 0.08 vs 1.28 ± 0.08, p<0.05) and rM1 (1.66 ± 0.09 vs. 1.40 ± 0.07, p<0.05). No significant difference was observed in rS1 (1.78 ± 0.07 vs. 1.70 ± 0.09, p = 0.18) and rV2 (1.57 ± 0.09 vs. 1.45 ± 0.07, p =0.18). Furthermore, modafinil significantly decreases the offset signal on rM1 (−2.59 ± 0.14 vs. −2.81 ± 0.09, p<0.05), whereas rOb (−2.92 ± 0.11 vs. −2.90 ± 0.17, p=0.10), rS1 (−2.41 ± 0.14 vs. −2.42 ± 0.20, p=0.9), and rV2 (−3.21 ± 0.21 vs. −3.40 ± 0.20; p =0.41) were unchanged. In contrast, as exhibited in Figure 4c, modafinil did not alter either the oscillatory component of the gamma power or the gamma peak frequency. These results suggest that the modafinil enhancement of gamma power is partially attributable to an increase in baseline broadband power. In other words, increases in gamma-band power are primarily driven by changes in the aperiodic component of the spectrum, rather than by enhanced oscillatory activity. Finally, the significant increase in the HFO power induced by modafinil should also be attributed to aperiodic activity, as the model did not detect a pure oscillatory component in this frequency band.

**Fig. 4.**
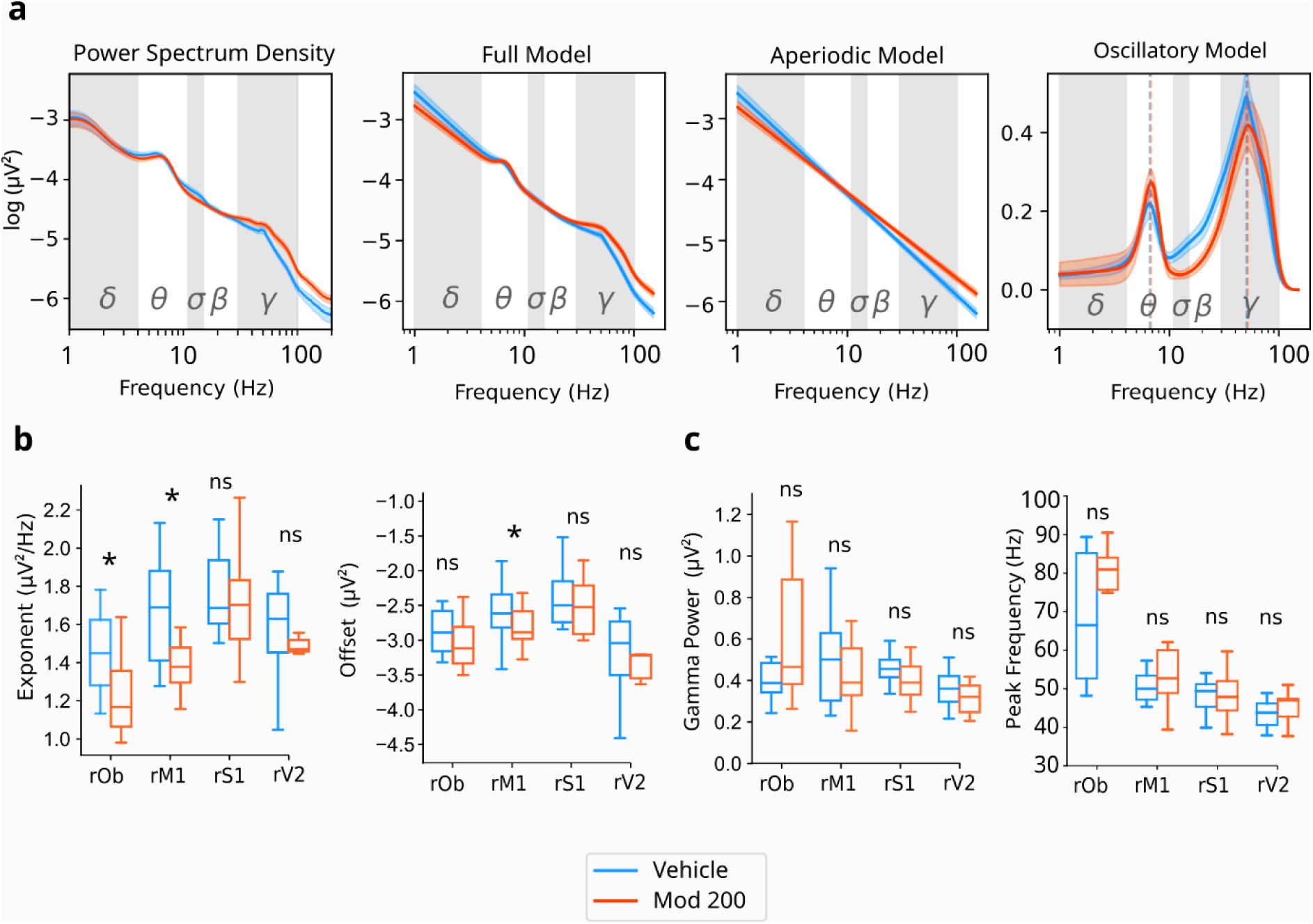
Modafinil modifies the aperiodic components of the spectral power during wakefulness. **(a)** Mean power spectrum and full model of FOOOF (aperiodic and periodic components) applied to rM1 cortex. **(b)** Comparison of aperiodic components (exponent at left, offset at right) between both conditions for each cortex. **(c)** Comparison of parametrized highest gamma aperiodic-adjusted power oscillation (left) and center frequency (right). Boxes represent the middle 50% of the data, stretching from the first quartile (Q1) to the third quartile (Q3). (*) indicates statistically significant difference (two-sided paired samples t-test) at p-value < 0.05. ns, not significant. rOb: right olfactory bulb (n = 6), rM1: right primary motor cortex (n=10), rS1, right primary somatosensory cortex (n=9), rV2: right secondary visual cortex (n=7)

### 3.3 Modafinil increased the power of slow waves during posterior sleep

Finally, we examined whether modafinil administration alters the spectral characteristics of subsequent sleep, despite not affecting its total hourly duration (see Figure 2). We first analyzed the time course of delta power across SWS. Figure 5a shows the relative delta power per hourly bin in the rM1 cortex across all states for vehicle and modafinil. Under vehicle conditions, relative delta power declined progressively over time, consistent with the typical homeostatic dissipation of slow-wave activity. In contrast, after modafinil administration, relative delta power increased, with the most pronounced differences observed during the 8th and 9th hour bins, which followed a sustained W (*p* < 0.05). To assess if the effect was generalized across all cortical brain regions, we analyzed the mean PSD of the delta band (0.5–4 Hz) post treatment during SWS for the 8th and 9th hour bin-combined. As shown in Figure 5b, modafinil significantly increased delta-band power in anterior regions, including the rOb (1.04 × 10^−3^ ± 2.30 × 10^−4^ vs. 6.60 × 10^−4^ ± 9.30 × 10^−4^ µV^2^/Hz, p < 0.05, n = 6), rM1 (2.18 × 10^−3^ ± 4.30 × 10^−4^ vs. 1.37 × 10^−3^ ± 1.90 × 10^−4^ µV^2^/Hz, p < 0.05, n = 8), and rS1 (3.08 × 10^−3^ ± 3.60 × 10^−4^ vs. 2.16 × 10^−3^ ± 3.40 × 10^−4^ µV^2^/Hz, p < 0.05, n = 7). In contrast, no significant change was detected in rV2 (7.50 × 10^−4^ ± 2.20 × 10^−4^ vs. 5.20 × 10^−4^ ± 1.50 × 10^−4^ µV^2^/Hz, p = 0.14, n = 6). No other frequency bands showed significant change when comparing modafinil and vehicle (see also supplementary Figure 3). Together, these findings indicate that subsequent sleep after modafinil-induced wakefulness is characterized by an increase in delta-band activity, particularly in anterior cortical regions.

**Fig. 5.**
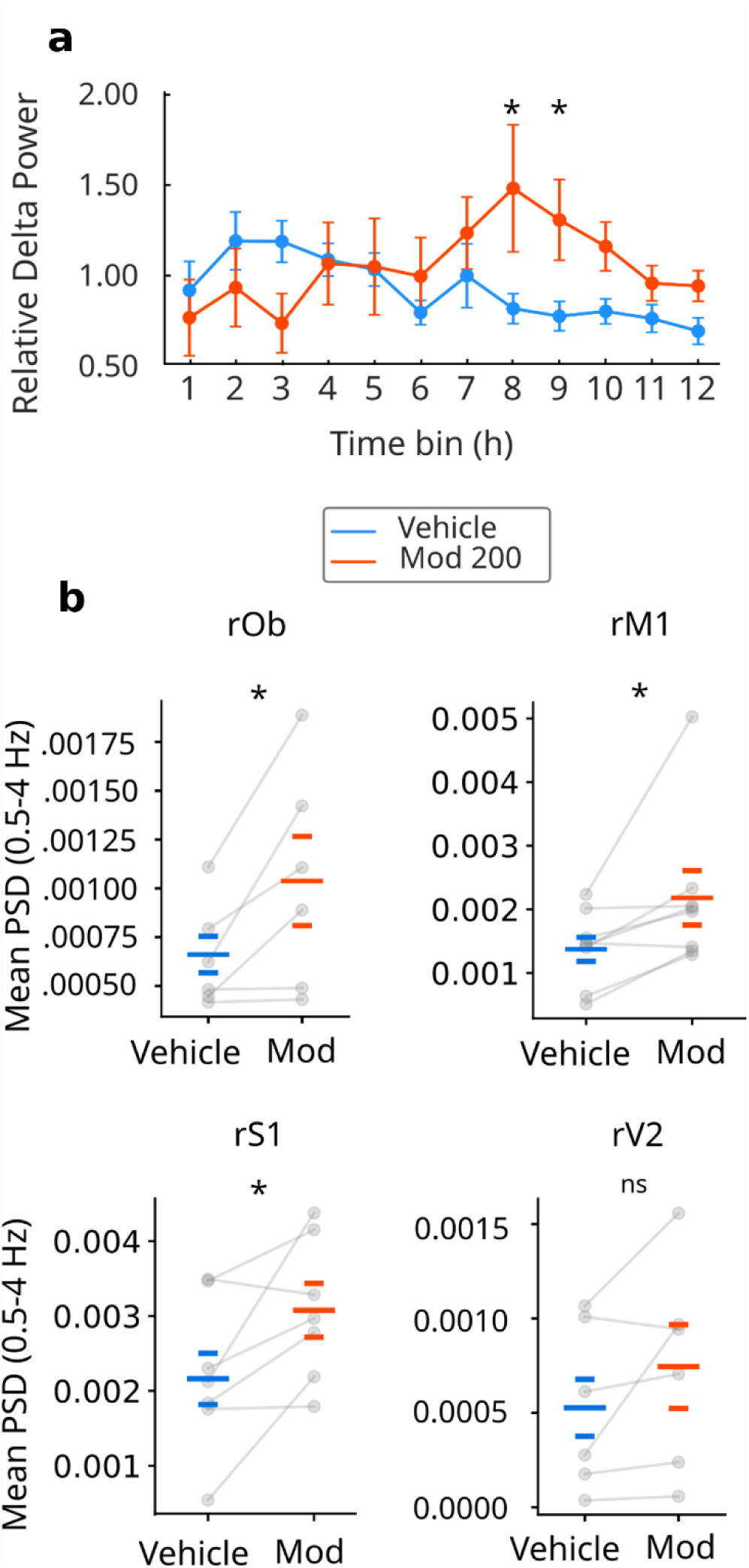
Effects of modafinil on delta power (0.5-4Hz). **(a)** Relative delta power (0.5–4 Hz) across hourly bins after vehicle or modafinil 200 mg/kg administration in the right primary motor cortex. Values are expressed as the mean delta power per hour in all behavioral states, normalized to the 12-hour mean delta power including all stages. Each dot represents the mean for that condition; error bars indicate SEM. (*) denote significant differences (two-sided paired t-test or Wilcoxon signed-rank test, p < 0.05). **(b)** Mean power spectral density of the SWS delta band between hours 8 and 9 after treatment, in different brain regions. Larger horizontal bars indicate group means, shorter bars indicate SEM. Lines connect paired values from the same animal. (rOb), right olfactory bulb; (rM1), right primary motor cortex; (rS1) right somatosensory cortex; and (rV2) right secondary visual cortex. (*) denotes significant differences (two-sided paired t-test, p < 0.05); ns, not significant

We also evaluated whether modafinil altered cortical oscillatory activity during REM sleep by computing the total PSD for all cortical sites. The analysis was performed during the 7–9 h post-treatment window, when animals begin to transition into REM sleep following the modafinil-induced W (see data above). No significant differences were observed between modafinil and vehicle across frequency bands throughout the course of REM sleep (see Supplementary Figure 4). These findings suggest that the spectral composition of REM sleep remains largely unchanged following modafinil-induced W. LS and IS are transitional states whose EEG patterns combine features of both wakefulness and slow-wave sleep, making them less suitable for quantitative frequency analysis.

## 4. DISCUSSION

We aimed to characterize changes in ECoG spectral composition during modafinil-induced W and its influence on subsequent sleep ECoG dynamics. The principal findings were: a) A dose-dependent increase in W time and a decrease in the hypnogram LZC. b) Gamma activity increased, as shown by power spectral density computed from W epochs, especially in anterior cortical brain regions. These changes were not mediated by changes in oscillatory components (neither gamma aperiodic-adjusted power nor peak frequency changed significantly). On the contrary, modafinil modified the aperiodic components of the ECoG. c) Following modafinil-induced W, total sleep time per hour was not significantly different from vehicle, although delta power increased in anterior cortical regions.

### 4.1 Modafinil increases wakefulness and reduces hypnogram complexity

Our finding that modafinil administration significantly increases time spent in W is consistent with previous reports in rodents, which demonstrated dose-dependent increases in W without rebound hypersomnolence (Duteil *et al*., 1990; Edgar and Seidel, 1997). This increase in W time was accompanied by a higher number of wake episodes and a prolongation of their mean duration, indicating a stabilization of the wake state. Such stabilization reduces the frequency of transitions between W and sleep states. Consistent with this interpretation, hypnogram architecture, assessed using LZC, decreased with higher doses of modafinil, reflecting longer, more predictable periods of W. While LZC has been previously applied to local field potentials to quantify cortical signal complexity (Abásolo et al., 2015), it has not been fully explored in the context of hypnograms (Catanzariti et al., 2025), representing a novel aspect of the present study. A closer inspection of the hypnograms showed heterogeneous responses, suggesting inter-individual variability in the wake-promoting effects of modafinil. This variability may reflect differences in intrinsic sensitivity to the pharmacological action of modafinil, as well as variability in drug absorption related to gastric content at the time of administration, given that animals were housed with *ad libitum* access to food and water. In humans, food intake has been shown to delay modafinil absorption (Robertson and Hellriegel, 2003); however, comparable data in rodents are lacking. Additional sources of variability may include inter-animal variability in the homogeneity of the modafinil suspension, leading to differences in absorption, as well as differences in the sleep duration before light onset, which may induce differential effects on the homeostatic sleep pressure (Everson *et al*., 1989).

### 4.2 Modafinil increases the power of the ECoG gamma band

We found that after modafinil administration, gamma power increased, mainly in anterior cortical brain regions. This anterior predominance is consistent with reports of stronger modafinil effects in frontal cortical regions (Gozzi *et al*., 2012). Previous reports in mice have shown an increase in gamma power with acute administration of modafinil (Hasan *et al*., 2009; Vas *et al*., 2020), but other studies in rats have not shown a significant increase (Claudio *et al*., 2019). Notably, in the latter case, a narrower gamma range was analyzed compared to our protocol. Importantly, previous pharmacological studies of modafinil reported increases in gamma power without distinguishing whether these changes reflected genuine oscillatory rhythms or other sources such as spiking-related broadband activity (Ray and Maunsell, 2011). More recent methodological work on EEG emphasizes the importance of separating true oscillatory activity—presumably arising from neuronal rhythms—from the aperiodic component of the power spectrum, which is ubiquitously present and may have distinct physiological generators and interpretations (Donoghue, Schaworonkow and Voytek, 2022). Applying this approach, our work shows that the increase in gamma power is mostly due to the aperiodic component. Previous studies suggest that such changes in the aperiodic component may reflect alterations in the excitatory–inhibitory balance (Gao, Peterson and Voytek, 2017), which is consistent with the described properties of modafinil as activating, wake-promoting, and enhancer of electrical coupling. Biophysical modeling of neurons shows that aperiodic signals can be influenced by neuronal and synaptic properties (such as activation of AMPA currents, GABA currents, and membrane properties) (Brake *et al*., 2024), and there is evidence that modafinil administration increases extracellular glutamate and decreases extracellular GABA, whose regional effects are dose-dependent and appear to be indirect (Tanganelli *et al*., 1992; Ferraro *et al*., 1997; Gerrard and Malcolm, 2007). The only neurotransmitter with evidence to be directly affected by modafinil is dopamine, probably through direct binding to dopamine transporter (Mignot *et al*., 1994; Wisor *et al*., 2001; Minzenberg and Carter, 2008). In this sense, an increase in dopaminergic transmission has been linked to an increase in gamma power. For example, evidence suggests an increase in gamma power following the administration of a dopaminergic agonist in hippocampal slices (Wang *et al*., 2022). In the same way, disease related to dopamine, like Parkinson’s disease, shows alterations in gamma synchrony related to dopamine levels (Lofredi *et al*., 2018), suggesting a link between gamma and dopamine. In this regard, selective experimental lesions of substantia nigra dopaminergic neurons result in decreased gamma-band ECoG activity in rats (Cavelli *et al*., 2019). One possible explanation for dopamine’s influence on gamma oscillations may depend on the fine tuning of the plasticity of parvalbumin basket cells throughout the modulation of their synaptic properties and electrical coupling (Bartos *et al*., 2002). These interneurons play a critical role in generating and maintaining gamma oscillation (Buzsáki and Wang, 2012). Even more, previous studies propose a role for specific dopamine receptors, like D4 receptors, on the gamma power (Andersson *et al*., 2012; Karunakaran *et al*., 2016). However, although D2 antagonists do not appear to affect the aperiodic component of the cat EEG (Gallo *et al*., 2024), the modulatory role of dopamine in shaping aperiodic activity has not yet been fully explored.

The gamma band is disrupted in cognitive disorders because of its broad involvement in cognitive functions, such as attention, long-term memory, working memory, and motor tasks; accordingly, both pharmacological and non-pharmacological approaches are beginning to be explored to target gamma oscillations for cognitive enhancement (Palacino, Manganotti and Benussi, 2025). If these cognitive disorders, as well as cognitive tasks, affect genuine oscillatory or aperiodic components, it remains under discussion. Interestingly, aperiodic components correlate with improved performance in cognitive tasks such as attention (Waschke *et al*., 2021); therefore, the spectral pattern we observe in our work is consistent with modafinil’s cognitive-enhancing profile, even though we cannot directly infer cognition due to our study limitations.

### 4.3 Modafinil effects on subsequent sleep

Classical sleep-deprivation studies established that sleep is governed by a homeostatic process sensitive to prior wakefulness, leading to compensatory changes in both sleep duration and sleep intensity (Borbély, 1982). Hence, the time spent in SWS and the amplitude/power of slow waves increase following sleep deprivation, providing a reliable marker of the sleep pressure accumulated during the deprivation protocol (Tobler and Borbély, 1986; Vyazovskiy *et al*., 2007). In line with earlier observations in animal models (Touret, Sallanon-Moulin and Jouvet, 1995; Edgar and Seidel, 1997; Lin *et al*., 2000), modafinil administration was not followed by a rebound in SWS duration episodes. However, an increase in delta power during this state was evident, indicating an accumulation of homeostatic sleep pressure during modafinil-induced wakefulness. In this sense, a study in mice has shown that delta power rebound after modafinil sleep deprivation is comparable to a non-pharmacological sleep deprivation (Kopp *et al*., 2002). Moreover, consistent with our findings, this study also reported a more pronounced increase in slow-wave activity over frontal cortices during sleep following modafinil-induced W. This regional difference might reflect greater functional activity of these regions during sustained W, as suggested by the spectral changes mainly seen in anterior regions. The cellular mechanism underlying this spectral change remains to be elucidated.

Although modafinil is commonly classified as a psychostimulant, its effects on sleep-wake architecture differ markedly from those of classical monoaminergic stimulants, which typically induce pronounced sleep disruption and rebound (Chapotot *et al*., 2003). Unlike caffeine (which promotes W by antagonizing adenosine A2A receptors and therefore intensifies the homeostatic sleep-pressure signal), modafinil does not interact directly with adenosine receptors (Urry and Landolt, 2015). Since the primary pharmacological effects of modafinil are mediated through dopaminergic signaling, one possible explanation for these findings is that dopamine and adenosine converge antagonistically within the same neuronal populations (Wisor, 2019). This antagonistic arrangement suggests that increased dopaminergic tone during modafinil-induced wakefulness could transiently counterbalance adenosine-driven sleep pressure. When dopaminergic drive diminishes after drug clearance, the unopposed adenosinergic influence may contribute to the subsequent enhancement of SWS. One potential site for such integrative interactions is the nucleus accumbens. Lesion studies indicate that the arousal-promoting effects of modafinil depend on the integrity of the nucleus accumbens core, but not the shell (Qiu *et al*., 2012). In parallel, adenosine A2A receptors are prominently expressed within the nucleus accumbens shell (Lazarus *et al*., 2011), where they are positioned to interact antagonistically with dopaminergic signaling, raising the possibility that the nucleus accumbens may serve as a hub where a modulation of the sleep-wake cycle may take place.

### Technical considerations

Technical considerations should be acknowledged. Not all the animals completed the 12-hour recording sessions, and in some cases, the ECoG data of some regions were incomplete. To mitigate this, we established a minimum of six animals to analyze each cortical region. Another aspect to consider is that gamma activity was measured in basal conditions, animals were not engaged in a specific cognitive task. Although we observed changes in the gamma band, these effects might be more pronounced or different when animals perform attention-demanding tasks under modafinil (Fries *et al*., 2001; Morgan *et al*., 2007). However, evidence from animals and humans suggests that the cognitive and attentional benefits of modafinil are more apparent under conditions of cognitive impairment, such as sleep deprivation, but not in normal conditions (Waters *et al*., 2005; Kredlow *et al*., 2019).

## 5. CONCLUSION

Our findings support that modafinil consolidates W in rats, decreases the temporal complexity of the sleep–wake cycle, and alters cortical dynamics by enhancing gamma-band power through changes in aperiodic rather than oscillatory components. Importantly, the wake-promoting effects of modafinil were not followed by increased SWS, but by a selective rise in delta power during subsequent sleep, indicating an influence on the intensity rather than the amount of sleep.

These results highlight the importance of distinguishing between oscillatory and aperiodic activity when assessing drug effects on brain rhythms, and they suggest that modafinil’s wake-promoting and pro-cognitive actions may be partly mediated through modulation of excitatory–inhibitory balance reflected in the aperiodic EEG signal. Future work should address how these cortical changes interact with behavioral and cognitive outcomes, and whether similar mechanisms underlie modafinil’s effects in sleep-deprived or clinical populations.

## Acknowledgements

This research was supported by the CSIC-I+D grupos 2022-group ID-22620220100148 grant, the “Programa de Desarrollo de Ciencias Básicas”, and “Agencia nacional de investigación e Innovación” from Uruguay.

## 7. STATEMENTS & DECLARATIONS

### 7.1 Funding

This work was supported by the Comisión Sectorial de Investigación Científica (Grant CSIC-I+D 2022-group ID-22620220100148) and by the Programa de Desarrollo de Ciencias Básicas (PEDECIBA) from Universidad de la República, Uruguay. Author M.M has received research fellowship from the Agencia Nacional de Investigación e Innovación (ANII) from Uruguay.

### 7.2 Competing Interests

The authors have no competing interests to declare that are relevant to the content of this manuscript.

### 7.3 Authors Contributions

Conceptualization: A.C., P.T., and M.M.

Methodology: M.M., J.P.C., and A.C.

Formal analysis and investigation: M.M. and D.M.M.

Writing-first draft of the manuscript: M.M.

Writing-review and editing: M.M., D.M.M., J.P.C., V.B., F.J.U., P.T., and A.C.

Funding acquisition: P.T. and A.C.

Resources: P.T. and A.C. Supervision: A.C. and P.T.

All authors have read and approved the final manuscript.

### 7.4 Data Availability

The datasets generated and analysed during the current study are available from the corresponding author upon reasonable request.

### 7.5 Ethics Approval

All experimental procedures performed in this study were conducted in agreement with the National Animal Care Law (#18611) and were approved by the Institutional Animal Care Committee (Comisión Honoraria de Experimentación Animal de la Universidad de la República, and the Ethics Committee of the Facultad de Medicina de la Universidad de la República, Uruguay (file N° 070151-000011-22)).

## Notes

### Competing Interest Statement

The authors have declared no competing interest.

