## Supplementary information for "Modafinil Modulation of Oscillatory and Aperiodic Brain Activity in Rats Across Wakefulness and Subsequent Sleep"

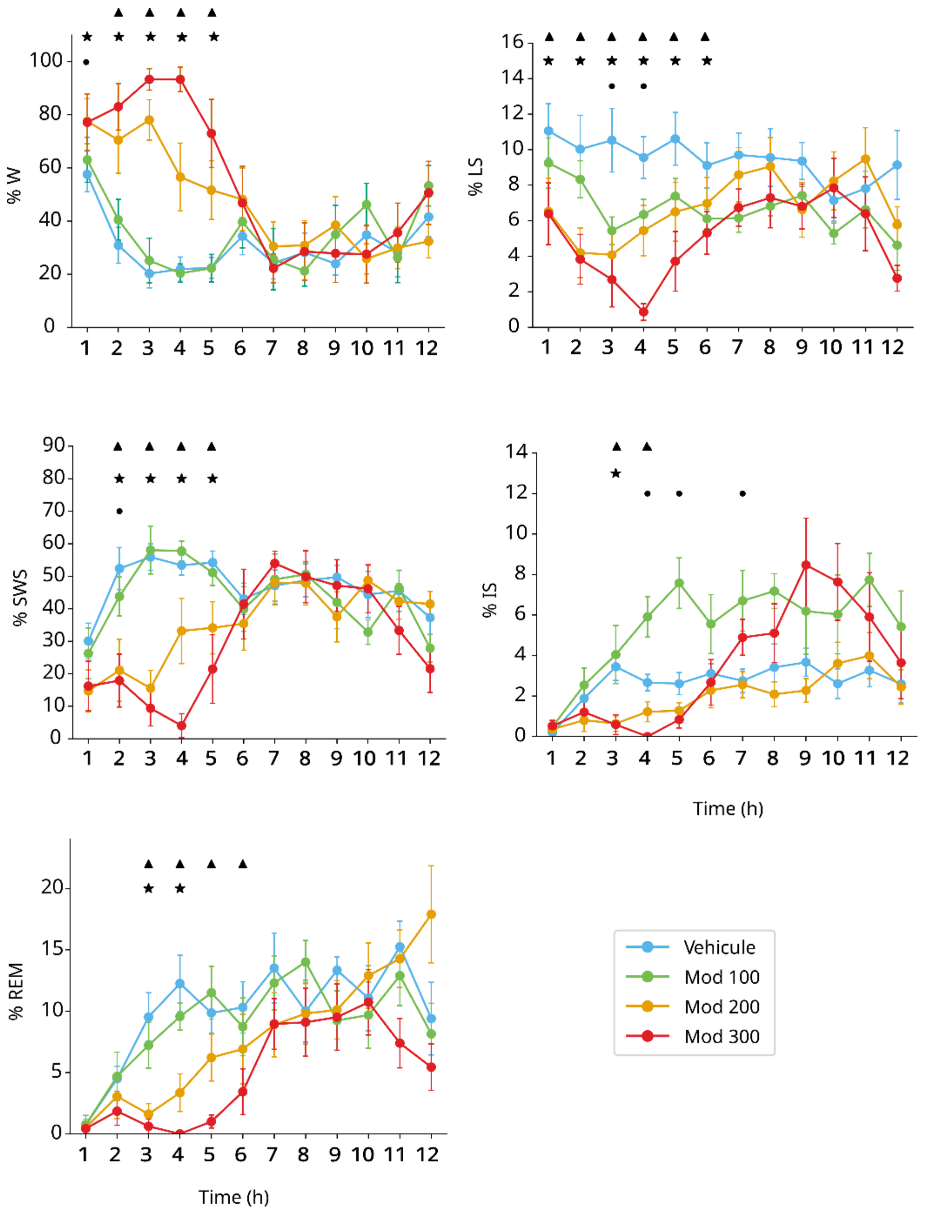


**Supplementary Fig. 1** Time course of the percentage of time spent in each state per hour (mean ± SEM), comparing vehicle vs modafinil 100 mg/kg, 200 mg/kg, and 300 mg/kg. A paired t-test was used to assess differences between vehicle and the different modafinil doses; significant differences ($p < 0.05$) are indicated by circles for vehicle vs. 100 mg/kg, stars for vehicle vs. 200 mg/kg, and triangles for vehicle vs modafinil 300 mg/kg. Sample size: n = 10 for hours 0 to 8; n = 7 for hours 9 to 12. For the 200 mg/kg comparison, the sample size was $n = 10$ for hours 0–8 and $n = 7$ for hours 9–12. For the 100 mg/kg condition, n =7. For the 300 mg/kg condition, n = 8 for hours 0-9 and n=7 for hours 9-12

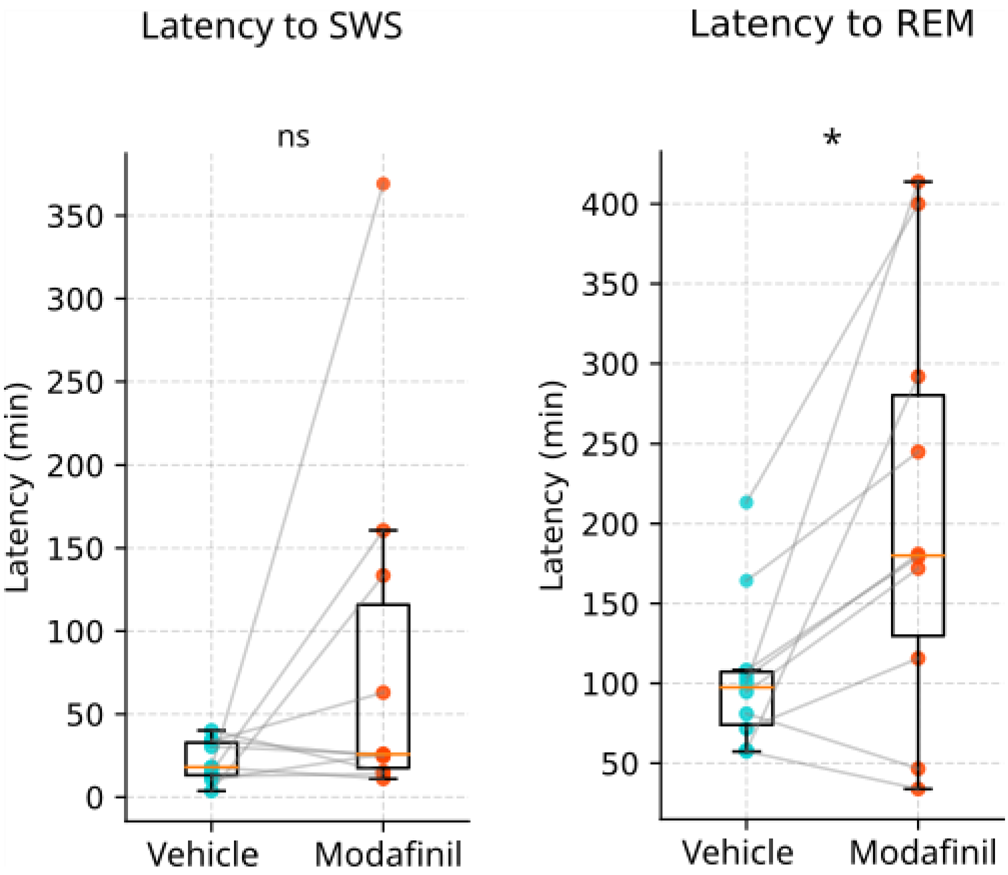


**Supplementary Fig. 2** Boxplots showing the latency to the first episode of slow-wave sleep (SWS) and rapid eye movement (REM) sleep following administration of vehicle or 200 mg/kg modafinil. Data are shown as median and interquartile range (Q1-Q3), with individual paired observations. A paired t-test revealed no significant effect of modafinil on SWS latency ($p = 0.12$; ns, not-significant), whereas REM sleep latency was significantly increased ($p < 0.05$ (*))

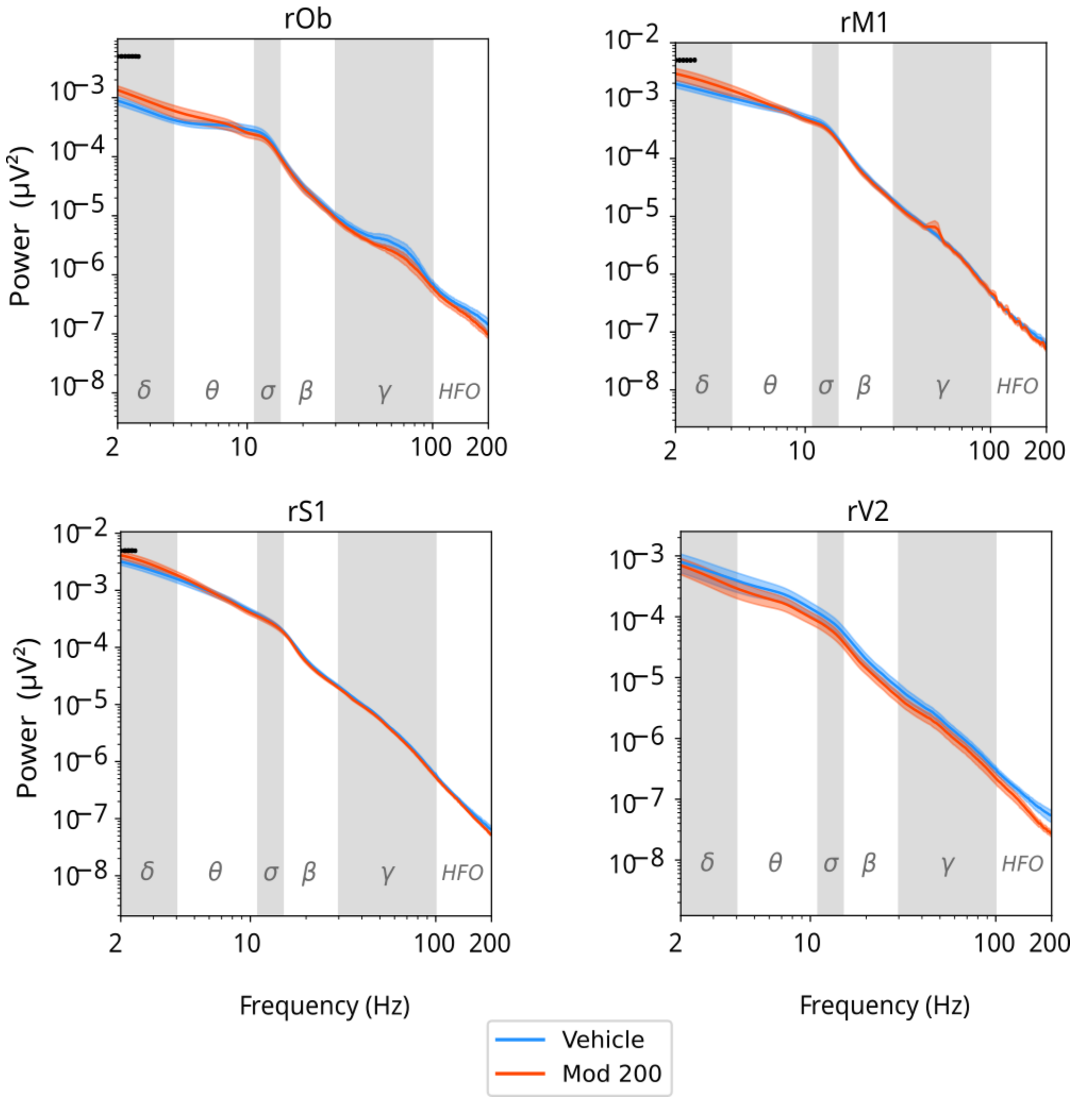


**Supplementary Fig. 3** Power spectrum density during slow wave sleep from hours 7 to 9 post-administration of all the analyzed cortices. Mean ± SEM. Background shading indicates canonical frequency bands: delta (0.5 - 4 Hz); theta (4 - 10 Hz); sigma (10 - 16 Hz); beta (16-30 Hz); gamma (30-100 Hz); and high-frequency oscillations (HFO, 100-200 Hz). Frequency bins showing significant differences between conditions (paired Student's t-test, $p < 0.05$) are marked with black dots. rOb: right olfactory bulb (n = 6), rM1: right primary motor cortex (n=8), rS1, right primary somatosensory cortex (n=7), rV2: right secondary visual cortex (n=6)

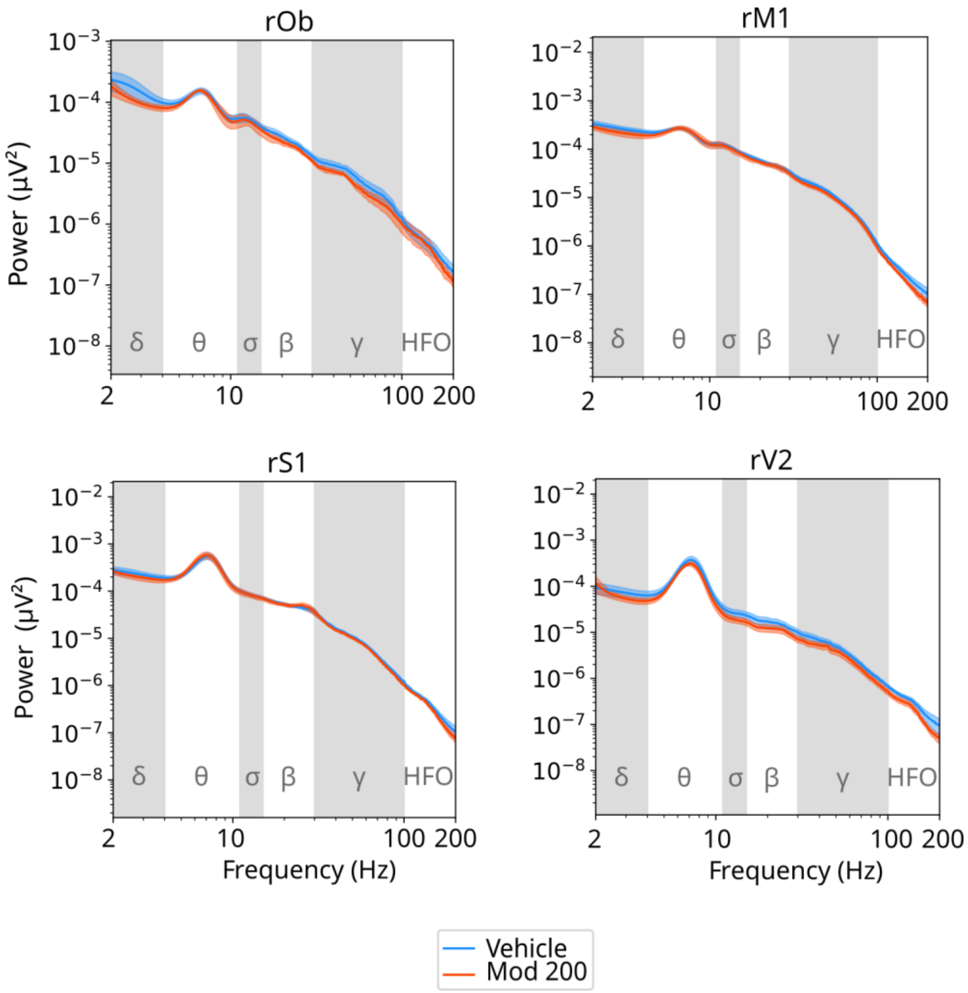


**Supplementary Fig. 4** Power spectrum density during REM from hours 7 to 9 post-administration of all the analyzed cortices. Mean ± SEM. Background shading indicates canonical frequency bands: delta (0.5 - 4 Hz); theta (4 - 10 Hz); sigma (10 - 16 Hz); beta (16-30 Hz); gamma (30-100 Hz); and high-frequency oscillations (HFO, 100-200 Hz). No significant differences between conditions (paired Student's t-test, $p < 0.05$) were detected. rOb: right olfactory bulb (n = 6), rM1: right primary motor cortex (n=8), rS1, right primary somatosensory cortex (n=7), rV2: right secondary visual cortex (n=6)
